# Leaf stage and roasting shape methylxanthine levels, chlorogenic acid, and overall metabolic profile of *Ilex vomitoria* leaf extracts

**DOI:** 10.1101/2025.09.29.679294

**Authors:** Ben Long, Scott Harding, Khadijeh Mozaffari, Jeffrey L. Bennetzen

**Author notes:** **Corresponding Author:** Ben Long https://github.com/bjl31194.

## Abstract

Yaupon holly (*Ilex vomitoria*) is a shrub native to the southeastern US whose leaves can be roasted and brewed into a caffeinated infusion. While production and consumption of yaupon tea is growing, not much is known about what influences downstream levels of caffeine and other metabolites within the leaf. To address this question, we used ultra-high performance liquid chromatography-mass spectrometry to measure the effect of leaf stage and roasting on the metabolome of yaupon leaves. We found caffeine levels strongly decreased with leaf stage but were not significantly affected by roasting. We also tentatively identified 100 chemicals that were dramatically increased by roasting, including several lactones and probable Maillard products related to known flavor compounds. These findings will help the yaupon tea industry produce a more consistent and tailored product, as well as lend insight into how leaf biochemistry changes over time.

## 1. Introduction

Faced with climate change and the sustainability issues surrounding coffee and tea plantations, it is vital to develop alternative sources of caffeinated beverages that can be grown in a wider range of environmental conditions (Davis Aaron et al., 2019; O’Brien Timothy & Kinnaird Margaret, 2003). Yaupon holly (*Ilex vomitoria* Ait., Aquifoliaceae) is a hardy shrub native to the southeastern United States that is a close relative of yerba maté (*Ilex paraguariensis*) (Yao et al., 2021). It is widely grown both as an ornamental and for its leaves, which can be brewed into a pleasant, caffeinated beverage (Power & Chesnut, 1919). Historically brewed by many Indigenous tribes in the American Southeast, yaupon tea has recently been subject to a resurgence of interest in wider circles as a sustainable alternative to tea and coffee (Hudson, 2004). A budding industry has since emerged in the southeastern US, comprised of a small but growing group of commercial producers.

Efforts to develop a yaupon tea industry have been bolstered by preliminary studies that found infusions made from the leaves and twigs are high in antioxidants and contain not only caffeine but the related bioactive methylxanthines theobromine and theacrine (Negrin et al., 2019; Palumbo et al., 2009). Another study found that polyphenols extracted from yaupon holly had anti-inflammatory and antioxidant effects in human colon cell culture, suggesting that consumption of yaupon tea may have nutraceutical benefits (Noratto et al., 2011). However, detailed work on the full chemical constituents of yaupon is lacking. There is also very little known about how levels of methylxanthines, antioxidants, and other compounds of interest change with regard to the stage of leaf harvested or post-harvest processing, even though these factors are taken under highest consideration in other beverage plants like *Camellia* tea (*C. sinensis*), coffee (*Coffea arabica* and *C. canephora*), and yerba mate (Aaqil et al., 2023; Gawron-Gzella et al., 2021; Wei & Tanokura, 2015).

In most modern commercial farms, yaupon leaves are usually harvested in bulk via hedge trimmer and frequently contain a mix of leaves from various developmental stages (Lou Thomann, personal communication, April 29, 2022). These leaves are dried in the dark at low humidity, then further processed. Post-harvest, there are two main processing methods. In the first, leaves are simply dried, crushed, and packaged to make a lighter product marketed as a “green” tea. The other method, which is more aligned with southeastern Indigenous tradition, involves roasting the dried leaves until they are visibly browned before crushing and packaging. A basic understanding of how leaf chemical composition changes — both regarding where leaves are harvested on the plant and whether or not they are roasted — could provide growers with the insight necessary to make a higher quality product with more nutraceutical value. This logic has been used to enhance the production of other beverage plants (Cai et al., 2022; Farag et al., 2022).

Roasting is a common method used to alter the flavor of many plant-based beverages, although the exact methods employed vary greatly. While coffee beans are always roasted to some degree, the process is optional in *Camellia* tea and yerba mate, with roasted variants mostly marketed as specialty products (Gawron-Gzella et al., 2021; Zhu et al., 2021). The heat from roasting catalyzes chemical reactions between green tissue components like sugars and organic acids and produces a wide array of new molecules that can have dramatic effects on sensory perception. These chemical changes are complex, including Maillard reactions, Strecker degradation of amino acids, and breakdown of other important compounds like chlorogenic acids, catechins, and glycosides (Farah et al., 2005; Wei & Tanokura, 2015; Zhu et al., 2021). There have been conflicting reports regarding how the roasting process affects methylxanthine levels, with some studies in coffee describing a decrease in caffeine with roasting and others reporting no effect (Awwad et al., 2021; Hečimović et al., 2011).

The stage of leaf at harvest is an important factor in the making of *Camellia* tea, with most product made from young leaves (Hagiwara & Wright, 2015). Chemical analyses have confirmed that indeed, methylxanthines are highest in these younger leaves, with levels decreasing in tissue harvested further down the stem (Song et al., 2012). This topic is less well-researched in yerba mate: Blum-Silva et al. (2015) found that leaf age significantly influenced methylxanthine levels in yerba mate leaves at 1, 2, and 6 months of age, but Dartora et al. (2011) reported no significant differences in methylxanthines or DPPH free radical-scavenging activity between 1 and 6 month-old leaf extracts (Blum-Silva et al., 2015; Dartora et al., 2011). Due to the highly variable nature of plant secondary metabolite production both within and across species, it is difficult to predict how the bioactive constituents of yaupon leaves will change with leaf age based on data from other species.

To answer these basic questions regarding leaf metabolites in yaupon, we harvested leaf tissue from three yaupon holly genotypes on three different stem locations, subjected them to either a green or roasted processing method, then conducted untargeted ultra-high performance liquid chromatography-mass spectrometry (UHPLC-MS) on the leaf extracts. We also quantified three methylxanthines (caffeine, theobromine, and theacrine) and a chlorogenic acid (3-caffeoylquinic acid).

## 2. Methods

### 2.1. Leaf collection and roasting treatment

Yaupon holly cuttings were collected at three different locations in Arkansas, Virginia, and Florida from wild plants in 2022-2023, rooted in a 1:1 mix of peat:vermiculite, and grown under greenhouse conditions for a year. Three genotypes of yaupon holly were used, each represented by three biological replicates (i.e., vegetative clones from the same mother plant). All plants were subjected to the same horticultural conditions: watered once a day, fertilized with a 15% nitrogen-9% phosphorous-12% potassium fertilizer every six months, and grown in full sun at a temperature of 21-26 °C. Overhead lights were set to a 13:11-hour day/night cycle. To minimize microclimatic variations within the greenhouse, individual plants were rotated and their positions on the bench shuffled every 7 days for the 3 months directly before harvest.

Leaf tissue was collected in April 2024 in the morning on a clear, sunny day. To measure the effect of leaf stage on metabolic composition, three classes of leaf were collected, each representing different stages of commercial harvest. “Young” leaves were defined as the first two leaves on an actively expanding shoot tip, usually lightest in color and the least waxy. “Softwood” leaves were taken from the 4^th^ and 5^th^ nodes down from an actively expanding shoot tip — these were usually darker and larger than young leaves and had an intermediate level of waxiness. Finally, “mature” leaves were taken from woody stems (i.e., past years’ growth). These leaves were always the darkest and most waxy. Leaf sizes ranged from 1.5-2.5 cm for young leaves and 2-3.5 cm for the softwood and mature stages.

Immediately following collection, leaves were transferred to a low-humidity chamber and dried at ambient temperature until they reached a constant weight, ∼4% w/w moisture content (Palumbo et al., 2007). Next, leaves from each biological replicate and stage were divided at the midrib and split into two treatments: green and roasted. For the roasting treatment, leaves were placed in an oven and roasted at 140 °C for 50 minutes, while green leaves were left at room temperature. Roasting temperature and time were determined by compiling information from multiple yaupon growers/roasters to best mimic conditions used in commercial production (Lou Thomann and Bryon White, personal communication, 2022). Following the treatment, all samples were ground to a fine powder using a bead mill (Tissuelyzer II, Qiagen). Samples were then sent to the Plant Metabolomics Laboratory at UGA for extraction and UHPLC-MS analysis.

### 2.2. Sample Extraction

Ten mg of dried, powdered leaf material was sonicated 2 x 15 min at 100 °C in 500 μl of 1:1 methanol:chloroform with internal standards ^13^C_6_ trans cinnamic acid and *d*_5_ benzoic acid. Then, 200 μl of HPLC-grade water was added, samples were vortexed, and then centrifuged for 5 min at 15,000 g at room temperature. The upper (aqueous) phase was passed through a 0.2μm PTFE filter (Agilent) and 1 μl of the filtered extract was injected for each analysis in positive or negative ion mode.

### 2.3. UHPLC-MS Analysis and Quantitation

Chemical diversity was assessed using an Agilent 1290 Infinity II ultra-high performance liquid chromatography module coupled to an Agilent 6546 quadrupole time-of-flight (QTOF) mass spectrometry (MS) detector with electrospray ionization probe and tandem MS (Agilent Technologies, Inc., Santa Clara, CA, USA). Liquid chromatography (LC) separations were carried out on an Agilent Zorbax RRHD C18 column with dimensions 2.1 x 50 mm and 1.8 μm particle size, maintained at 45 degrees C. The elution gradient (0.5 ml/min) started at 0%B for 0.5 min, went to 15%B at 3.5 min, to 30%B at 4.5 min, to 50%B at 5.5 min and then to 100%B at 7.0 min, with a 2 min hold at 100%B. Each run was followed by a 3 min column re-equilibration step. Solvent B was 100% acetonitrile with 0.1% formic acid. Solvent A was 100% HPLC grade water with 0.1% formic acid. MS conditions were as follows: nebulizer gas flow was 45 psi, drying gas temperature was 300 °C with a nebulizer sheath temperature of 250 °C, capillary voltage was 4000 V, nozzle voltage was 500 V, and the fragmentor voltage was 125 V. MS acquisition rate was 3 spectra/sec with a duration per spectrum of 0.33 sec. The data storage threshold value was 200 (absolute threshold and 0.01% for relative threshold). Mass-to-charge ratio ranged from 100-1700, and the instrument accuracy at the time of analysis was 10 ppm (parts per million) in the range of 50-650 m/z.

To monitor instrument performance and signal stability, QC samples of known concentration (e.g., 3 ppm of an external standard different from the analytes being quantified) were injected periodically throughout the run. Injection order was also randomized to minimize run-order effects and reduce systematic bias. While an external standard was not included, pooled QC samples, prepared by mixing aliquots from each biological sample, were injected after every 15 runs. These were used to monitor instrument performance, assess signal drift, and evaluate technical reproducibility. Reagent blanks were analyzed after every three injections to assess potential carryover or background contamination.

Preliminary testing revealed that under these conditions positive and negative ion mode could be used together to yield a larger set of metabolites than did either mode alone. In general, metabolites of keen interest including the methylxanthines ionized best using positive ion mode while phenylpropanoid containing substances like chlorogenic acids ionized best using negative ion mode. Therefore, two separate full-scan acquisitions were performed, one in negative ion mode and one in positive ion mode. Each was optimized separately to maximize sensitivity and data quality.

Compound IDs were obtained using the ID-browser function of MassProfiler v 11.0 (Agilent Technologies, Inc., Santa Clara, CA, USA). The Agilent METLIN database (v B.08.00) was the primary library used for compound identification, supplemented with the NIST20 Mass Spectral Library and in-house authentic standards for further confirmation. Features that exhibited strong spectral matches in public databases such as METLIN, supported by accurate mass and isotope pattern consistency but without confirmation by authentic standards, were classified as tentatively identified. Features were classified as identified compounds if they matched authentic standards based on retention time, MS/MS spectra, and accurate mass (within ±5 ppm). LC-MS grade standards for caffeine (Chromadex #ASB-00003032-010), theobromine (Chromadex #ASB-00020248-010), theacrine (Cayman Chemical #26855), and chlorogenic acid (3-CQA, Chromadex #ASB-00003450-005) were purchased and used to generate calibration curves for quantitation (Chromadex under Niagen Bioscience, Los Angeles, CA, USA; Cayman Chemical Company, Ann Arbor, MI, USA). Curves exhibited good linearity (R^2^ > 0.97) across the range of quantified metabolites. Calibration curves used 1/x weighting to improve accuracy at low concentrations. Isotopically labeled internal standards were also included to correct for matrix effects and variability. Limits of detection for all three methylxanthines and 3-CQA were 0.005 ng/ul.

### 2.4. Data filtering and statistical analysis

All analyses downstream from peak calling were performed in MetaboAnalyst 6.0 (Pang et al. 2024) and R (v4.4.3; R Core Team 2025). R scripts used are available at https://github.com/bjl31194/yaupon-metabolomics.git. Peak intensity and quantification data were normalized using ^13^C_6_ trans cinnamic acid and *d*_5_ benzoic acid internal standards. Metabolites were flagged as potential contaminants and discarded from the dataset if they were detected in the blank sample at levels higher than 1/3 the average value of the real samples. Finally, metabolites with missing values (likely below the detection threshold) were replaced with 1/5 of the minimum value found in the other samples.

For quantified metabolites, differences in the mean concentrations between the three leaf stages were tested for significance using a Kruskal-Wallis test followed by post-hoc Fisher’s LSD tests between individual stages. Significant differences in quantified metabolites between green and roasted treatments were assessed using Student’s *t*-tests.

To uncover other metabolites that increased or decreased due to roasting, *t*-tests were conducted on peak intensity data for the full set of metabolites between treatments. Raw *p*-values were corrected for multiple testing using the false discovery rate (FDR) method. Metabolites with an FDR-adjusted *p*-value less than 0.05 and a fold change between treatments greater than 2 or less than -2 were classified as increased or decreased by roasting, respectively.

Changes in metabolic composition with leaf stage were assessed using multiple methods. First, principal component analysis (PCA) was performed on the full set of samples and metabolites to check whether samples grouped more strongly by treatment or stage. Then, a multiple linear model was used to find metabolites that correlated strongly with stage. Kruskal-Wallis tests with an FDR-adjusted *p*-value to account for multiple testing were used to assess the significance of leaf stage differences in these correlated metabolites. Finally, random forest analysis was conducted to select metabolites/features important in classifying samples by stage. Metabolites that had an adjusted *p*-value < 0.05 and were ranked in the top 25% of variables in the random forest analysis were selected as being significantly associated with stage. Linear coefficient values and fold change between classes (with an absolute value > 2) was used to determine the direction of change (e.g., up with maturity or down with maturity).

## 3. Results

### 3.1. Methylxanthine and Chlorogenic Acid Quantitation

Four metabolites with known effects on beverage quality, namely the methylxanthines caffeine, theobromine, theacrine (three methylxanthines) and the polyphenol 3-CQA, were all detected in unroasted and roasted leaves. Caffeine and methylxanthine levels varied considerably between genotypes and leaf stage (Table 1).

**Table 1.** Standards and reagents used in the UHPLC-MS analysis.

| Reagent | Quality/Purity | CAS | Product/Supplier |
| --- | --- | --- | --- |
| Caffeine | Primary (HPLC reference standard-grade, $\geq 99.9\%$) | 58-08-2 | #ASB-00003032-010 (Niagen Bioscience, Los Angeles, CA, USA) |
| Theobromine | Primary (HPLC reference standard-grade, $\geq 99.9\%$) | 83-67-0 | #ASB-00020248-010 (Niagen Bioscience, Los Angeles, CA, USA) |
| Theacrine | Analytical reference standard grade, $\geq 98\%$ | 2309-49-1 | #26855 (Cayman Chemical Company, Ann Arbor, MI, USA) |
| Chlorogenic acid (3-CQA) | Primary (HPLC reference standard-grade, $\geq 95.5\%$) | 327-97-9 | #ASB-00003450-005 (Niagen Bioscience, Los Angeles, CA, USA) |
| Acetonitrile | UHPLC grade | 75-05-8 | Sigma-Aldrich 900667-4L (Burlington, MA, USA) |
| Water | UHPLC grade | 7732-18-5 | Sigma-Aldrich<br>900682-4L<br>(Burlington, MA,<br>USA) |

**Table 2.**
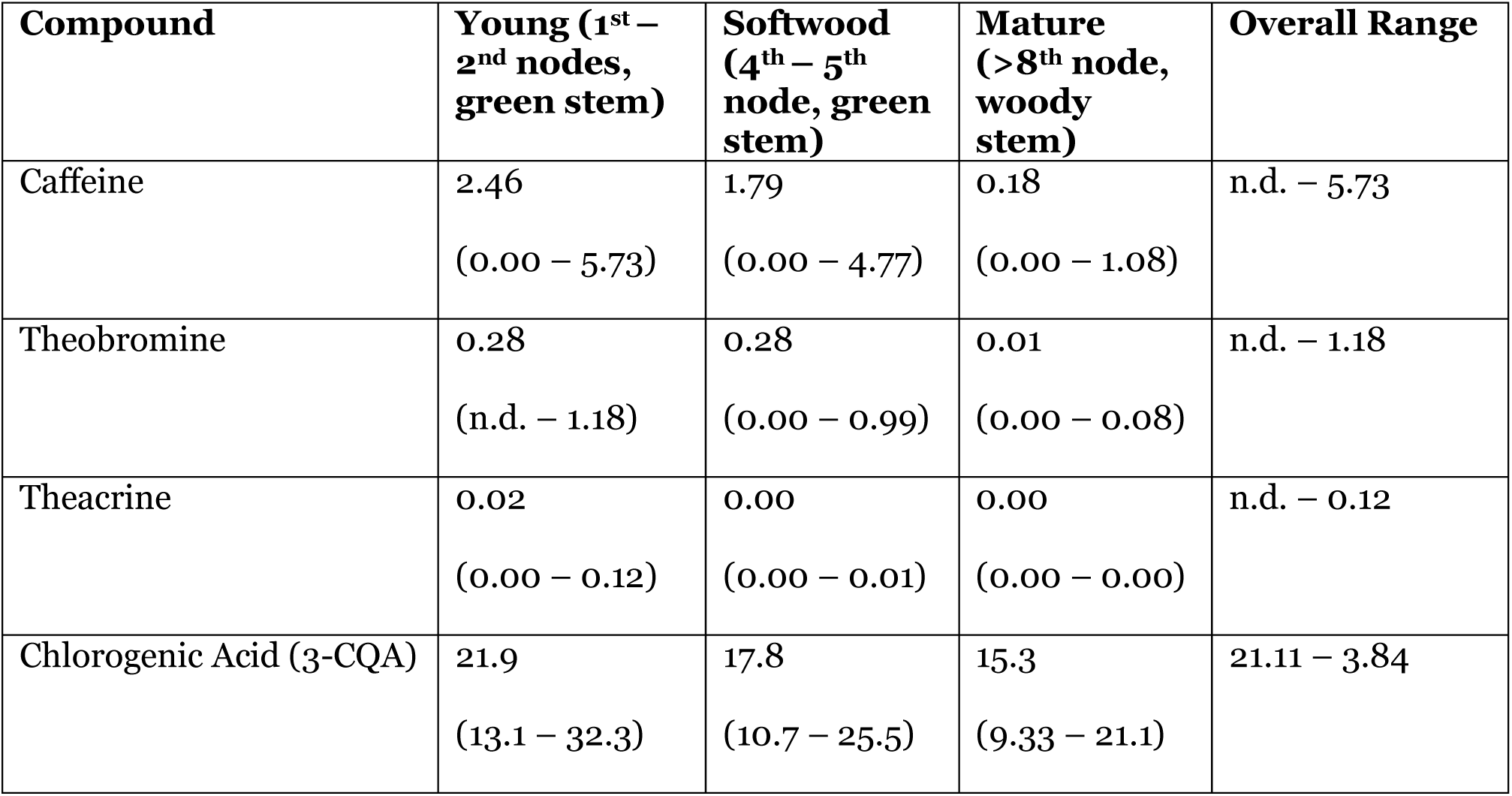
Levels of 3-caffeoylquinic acid and methylxanthines detected in yaupon leaves. Values are given in mg/g.

| Compound | Young (1<sup>st</sup> – 2<sup>nd</sup> nodes, green stem) | Softwood (4<sup>th</sup> – 5<sup>th</sup> node, green stem) | Mature (>8<sup>th</sup> node, woody stem) | Overall Range |
| --- | --- | --- | --- | --- |
| Caffeine | 2.46<br>(0.00 – 5.73) | 1.79<br>(0.00 – 4.77) | 0.18<br>(0.00 – 1.08) | n.d. – 5.73 |
| Theobromine | 0.28<br>(n.d. – 1.18) | 0.28<br>(0.00 – 0.99) | 0.01<br>(0.00 – 0.08) | n.d. – 1.18 |
| Theacrine | 0.02<br>(0.00 – 0.12) | 0.00<br>(0.00 – 0.01) | 0.00<br>(0.00 – 0.00) | n.d. – 0.12 |
| Chlorogenic Acid (3-CQA) | 21.9<br>(13.1 – 32.3) | 17.8<br>(10.7 – 25.5) | 15.3<br>(9.33 – 21.1) | 21.11 – 3.84 |

All four compounds had significant abundance associations with leaf stage (Fig 1). Caffeine, theobromine, theacrine, and 3-CQA maxima were highest in the youngest leaves, with maximum values decreasing markedly after the softwood leaf stage. These patterns were found regardless of genotype. Roasting did not have a significant effect on methylxanthine or 3-CQA content.

**Figure 1.**
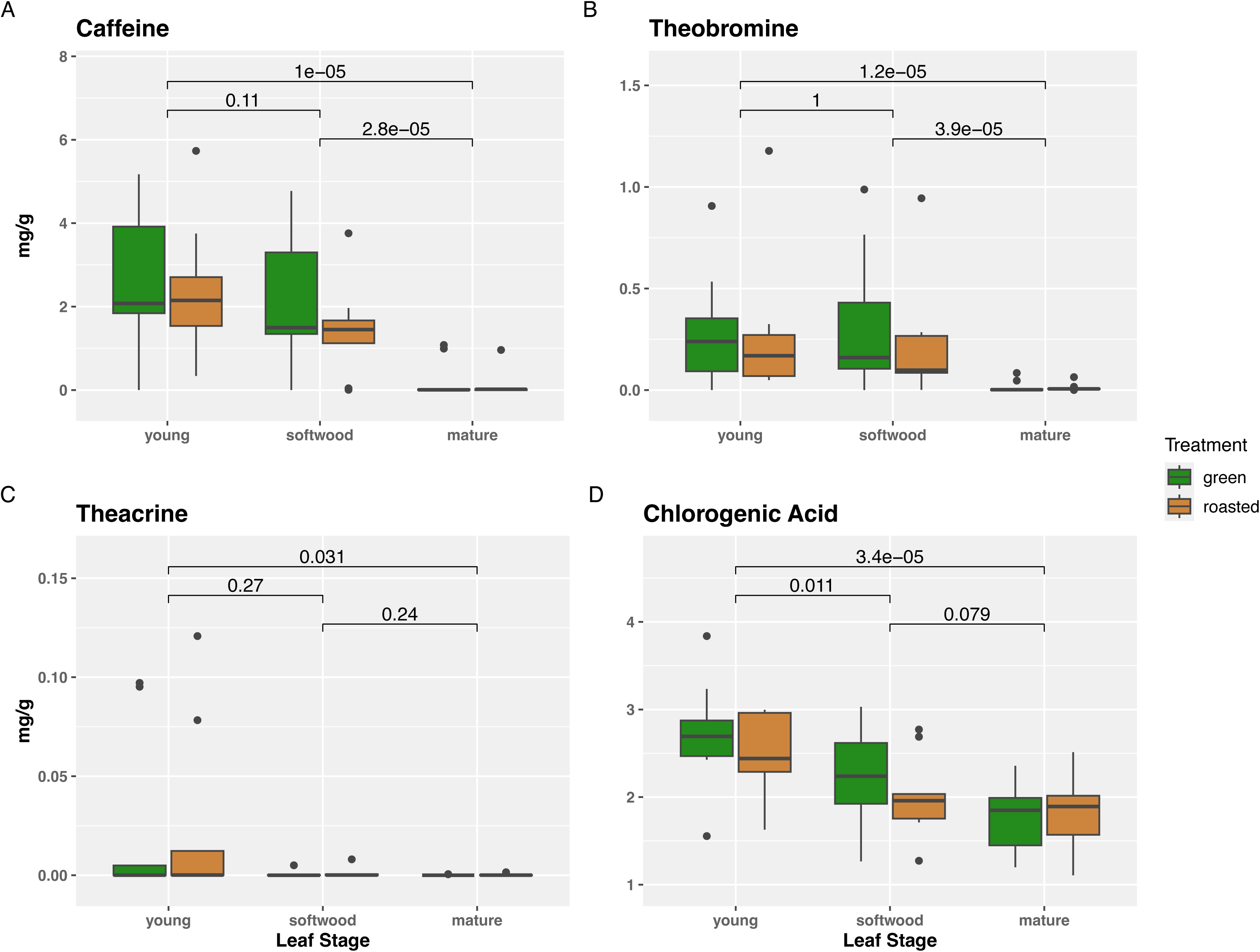
Methylxanthine and chlorogenic acid content of yaupon leaves across all genotypes, grouped by leaf harvest stage and roasting treatment. (A) Caffeine, (B) Theobromine, (C) Theacrine, and (D) Chlorogenic acid (3-caffeoylquinic acid). Numbers above brackets indicate p-values obtained from Fisher’s LSD post-hoc tests between groups.

### 3.2. Chemicals Affected by Roasting

Approximately 100 tentatively identified compounds were found to significantly increase with roasting: 74 detected in negative mode, 24 in positive mode, and 2 in both. Those most elevated included caffeoylshikimic acids, larixinic acid, triterpenoid saponins, lactones, quinones, and a ketal. Fewer chemicals decreased with roasting: 62 total, with 7 in negative mode, 54 in positive mode, and 1 detected in both (Fig 2). These included several glycosides and alkaloids, as well as multiple isomers of 3,4-dicaffeoyl-1,5-quinolactone. While the increase in certain chemicals was quite dramatic (up to 135-fold in roasted samples on average), PCA indicated these roasting-induced changes were not responsible for most of the variance between samples (Fig S1).

**Figure 2.**
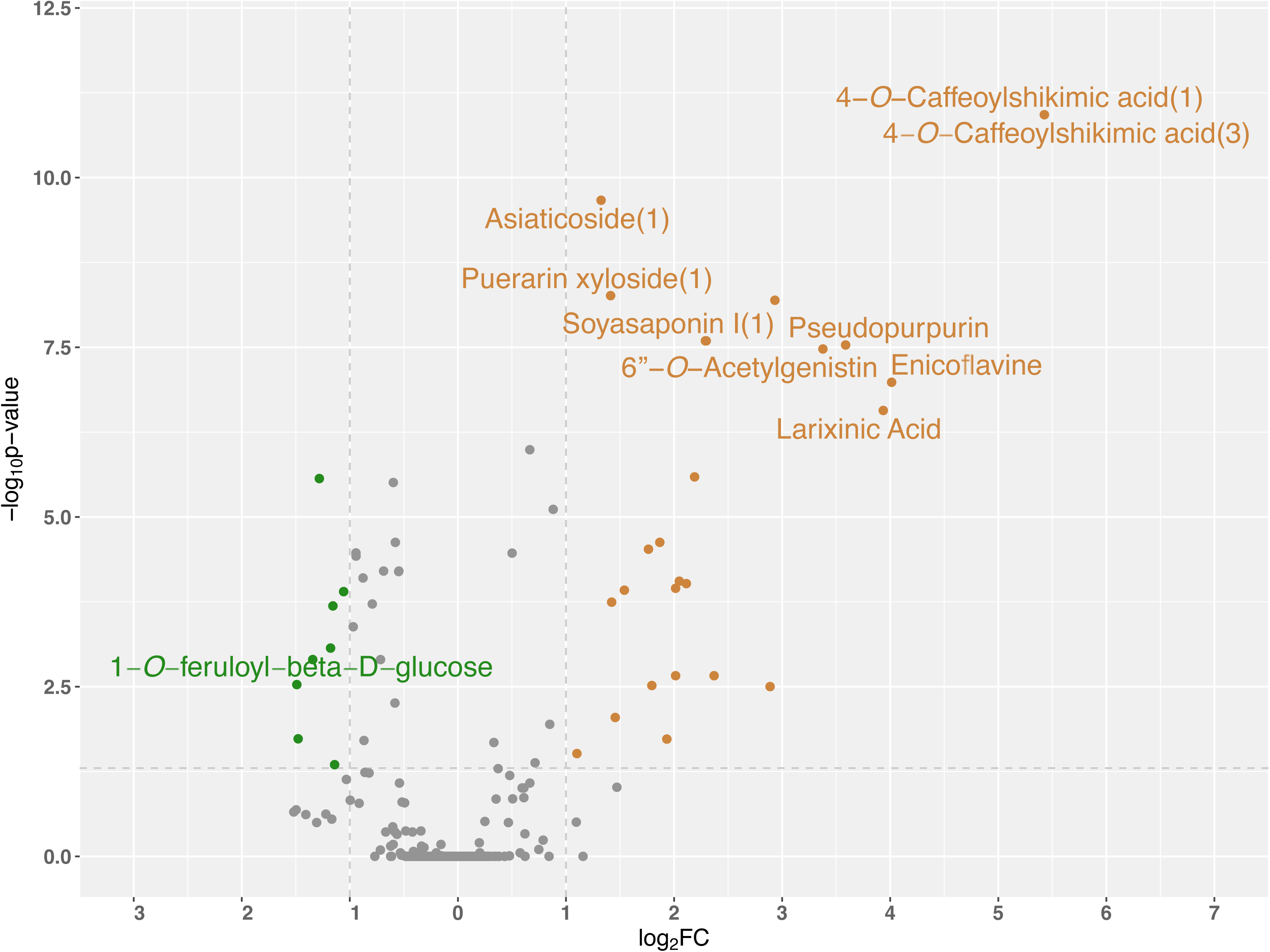
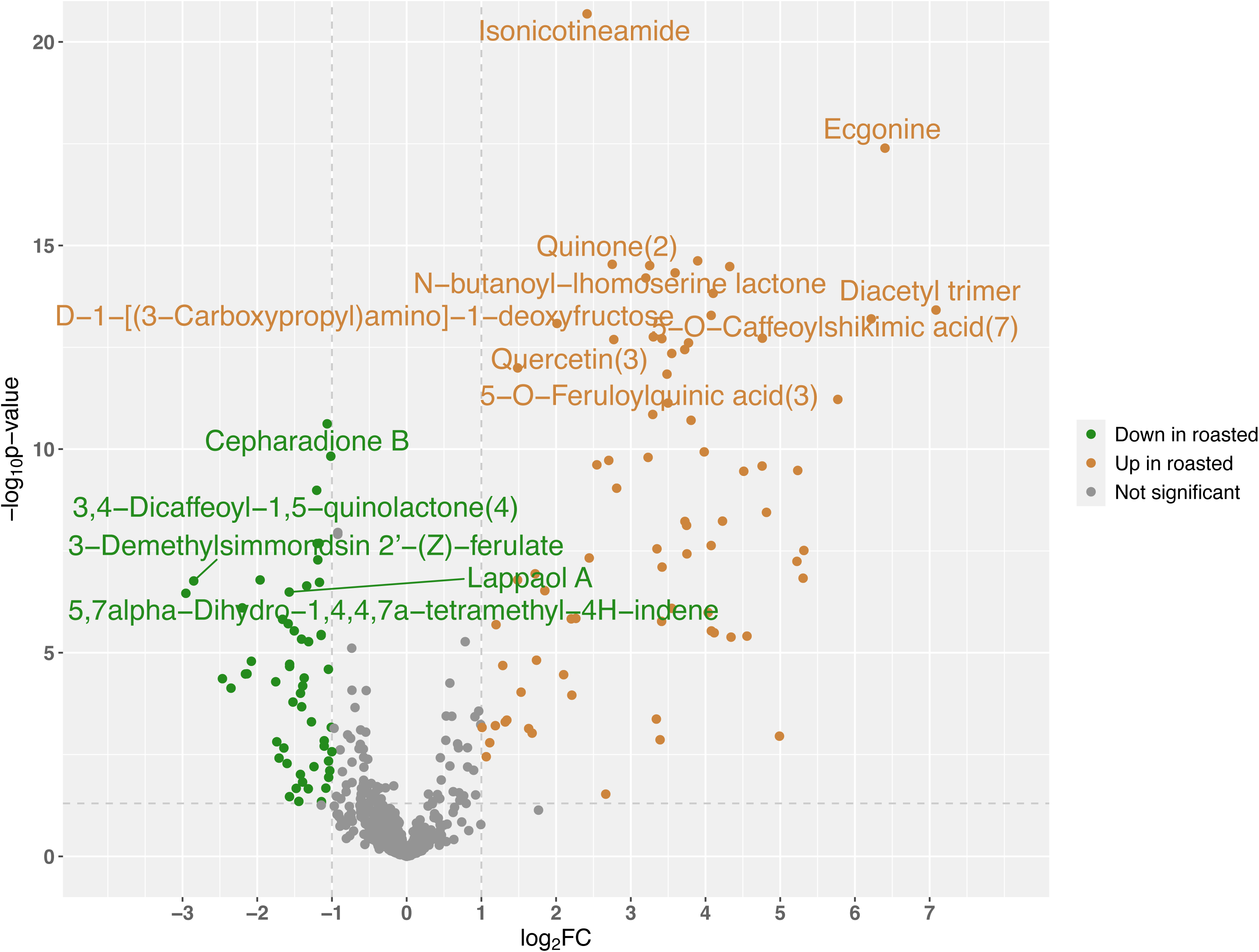
Compounds significantly affected by the roasting process. (A) Chemicals better detected in negative ion mode, (B) Chemicals better detected in positive ion mode. Compounds for which multiple isomers were detected were numbered according to the specific isomer; e.g., chlorogenic acid(1).

### 3.3. Metabolic changes associated with leaf stage

Overall, strong associations were found between metabolic composition and leaf developmental stage at harvest. Metabolite profiles from each of the three stages clustered together when visualized on a PCA scores plot, with the softwood group overlapping heavily with the young group (Fig 3). PC1 and PC2 were able to explain 45.2% and 19.8% of the variance between samples, respectively.

**Figure 3.**
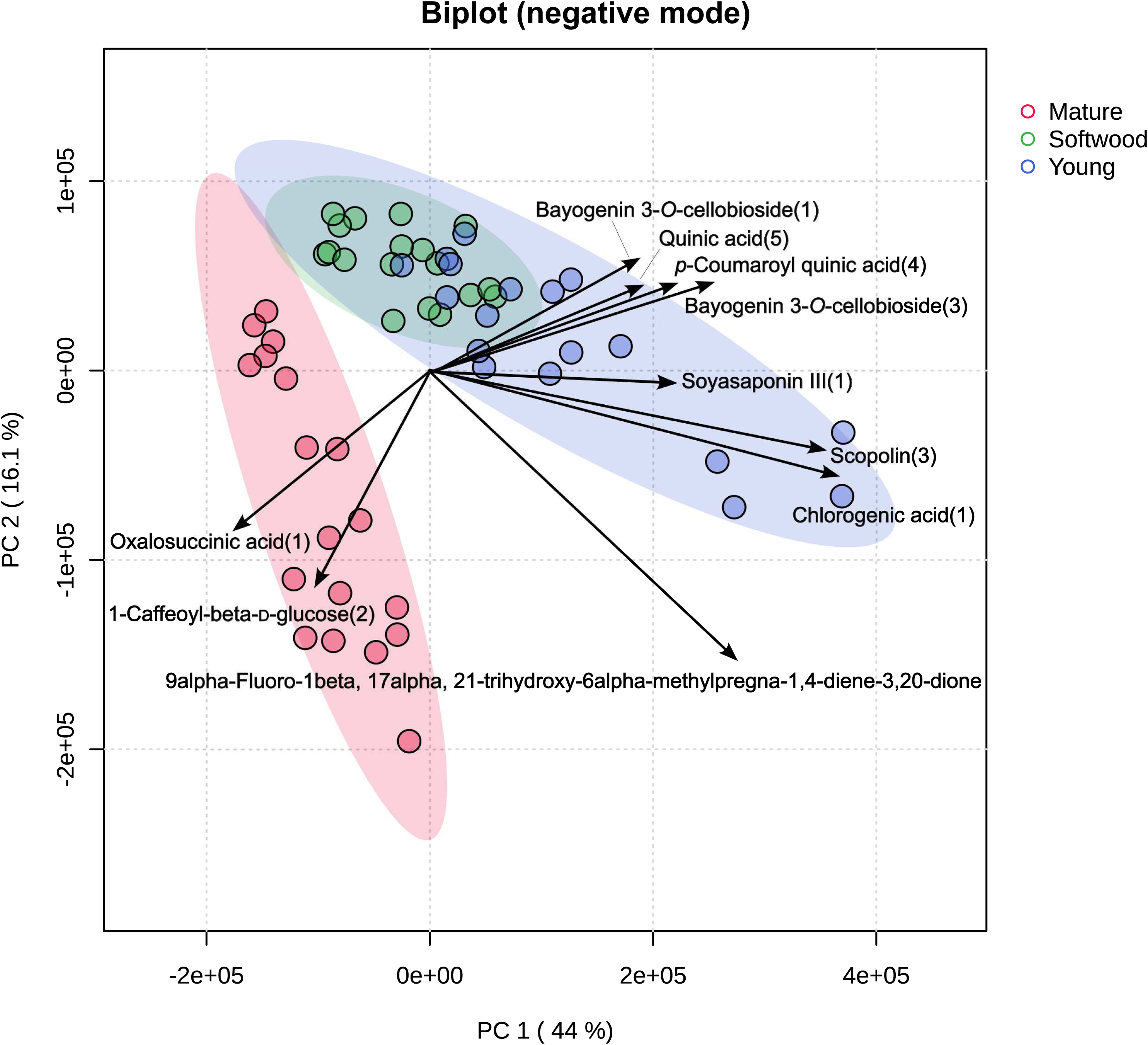
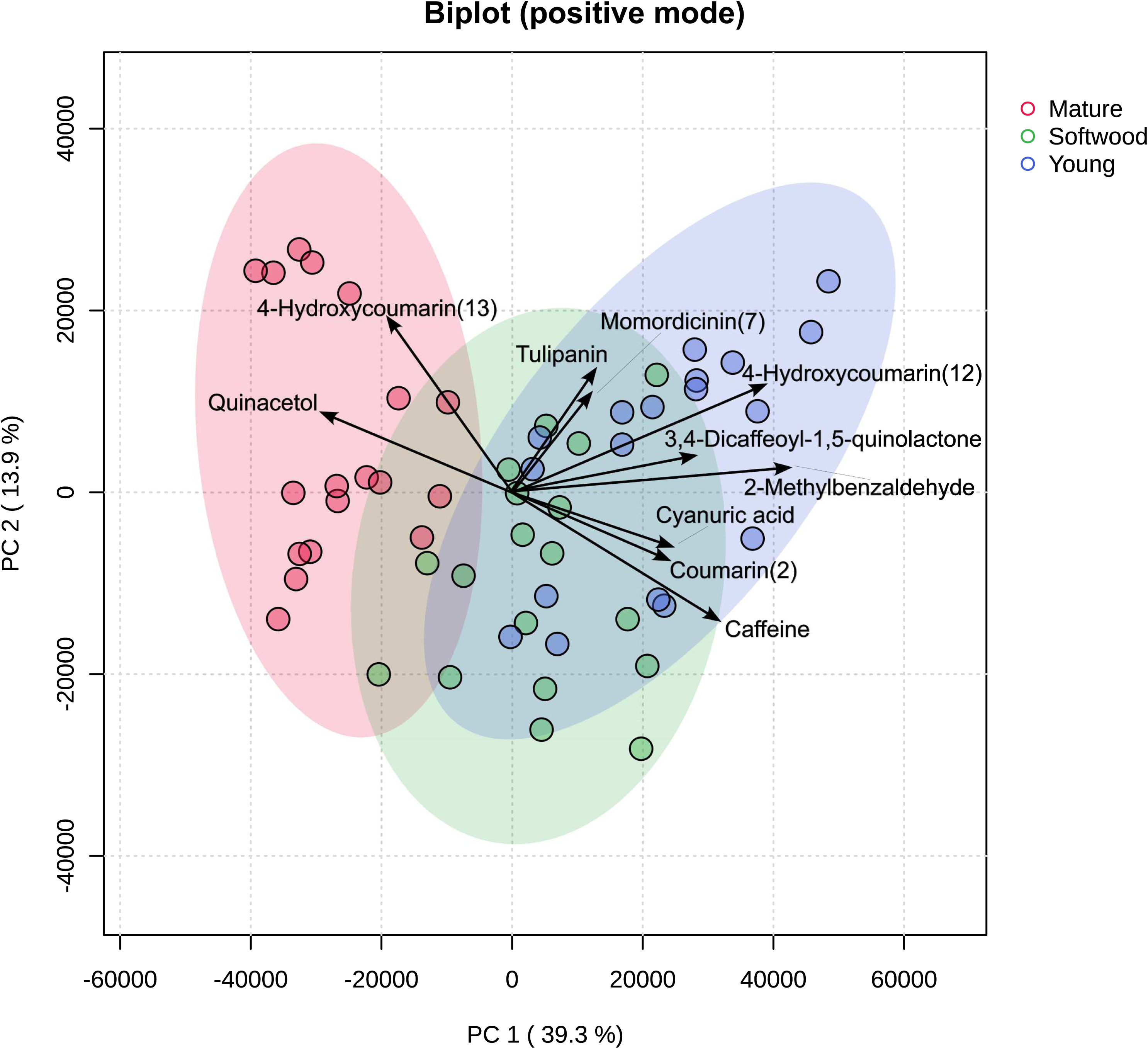
PCA biplot of samples run on UHPLC-MS, in negative and positive modes, grouped by leaf stage. (A) Negative mode results, (B) Positive mode results. Compounds for which multiple isomers were detected were numbered according to the specific isomer; e.g., chlorogenic acid(1).

Out of the 753 compounds tentatively identified in negative mode, we found 236 that were significantly correlated with leaf stage; e.g., significant in the linear model results and ranked highly in random forest analysis. A total of 96 of these were elevated in mature leaves, while 140 were depleted. From the 1141 compounds tentatively identified in positive mode, 139 were significantly elevated in mature leaves and 135 were depleted. These chemicals meeting the significance criteria are summarized in Tables S1 and S2, where those with a fold change greater than 2 (i.e., increase with age) are labeled “UP” and those with fold changes less than -2 (i.e., decrease with age) are labeled “DOWN.” Multiple isomers of some compounds, including chlorogenic acid, were detected.

## 4. Discussion

Here, we investigated the effects of leaf stage at harvest and roasting treatment on the metabolome of yaupon leaves, two factors studied in the context of other beverage plants but which have never been examined in yaupon holly.

The highest level of caffeine detected among the wild yaupon holly accessions examined in this study was 0.57% of dry weight, found in the youngest leaves. This is less than values reported in the literature for *Coffea arabica* (∼1.0%), young shoots of *Camellia sinensis* (∼2.8%), or yerba maté (∼1.0%) (Komes et al., 2009; Lean et al., 2011). The highest theacrine level detected was 0.12% of dry weight, also found in young leaves. This value is comparable to theobromine levels reported in commercial yerba maté (0.1-0.2%), although the young yaupon leaves examined here contained less theobromine when averaged across genotypes, 0.03% of dry weight (Lean et al., 2011; Negrin et al., 2019). Young yaupon leaves also contained theacrine, a bioactive methylxanthine derivative of caffeine that has been detected in wild coffee relatives and a few varieties of *Camellia*, most notably *C. assamica* var. kucha, but not in most *Arabica* coffee or common tea varieties (Ashihara et al., 2011; Zheng et al., 2002). Theacrine has become a compound of interest for human health, as some research has shown supplementation in humans improves mood, energy, and concentration (Ziegenfuss et al., 2017). However, theacrine levels in the accessions used in this study did not exceed 0.01%, much lower than values reported in young leaves of *C. sinensis* var. kucha (1.4% mg/g). Therefore, theacrine concentration may be an attractive target for future breeders interested in improving yaupon’s unique chemical composition compared to other caffeinated beverages. Finally, 3-CQA levels in young yaupon leaves (1.3-3.2% dry weight) were comparable to reported values in coffee and yerba mate (Butiuk et al., 2016; Farah et al., 2006). Put together, these findings suggest that yaupon could find a niche as a lower-caffeine alternative to other caffeinated beverages that is still high in antioxidant chlorogenic acids.

Both methylxanthine and 3-caffeoylquinic acid levels between samples were strongly correlated to leaf stage, with all four chemicals decreasing in leaves harvested further down the stem — regardless of genetic background. Small differences were found between young and softwood leaves, but the shift in methylxanthine content between mature leaves and the other two classes was dramatic. Indeed, many mature leaf samples did not contain detectable levels of methylxanthines at all. These results suggest that growers may be able to increase caffeine and antioxidant levels in their finished product by simply restricting harvest to the current year’s growth, as opposed to an even mix of leaf ages.

Our findings are also in line with similar research conducted in *Camellia* tea, where caffeine levels were found to decrease as leaf samples were taken further down the stem (Lu et al., 2009). This trend has also been observed in yerba mate in most (but not all) studies (Blum-Silva et al., 2015; Dartora et al., 2011; Esmelindro et al., 2004) Considering the documented insecticidal and antioxidant properties of methylxanthines and chlorogenic acids, respectively, this pattern also agrees with the hypothesis that younger leaves rely more on chemical defenses for protection than older ones: as leaves grow thicker and waxier, they become less attractive to herbivores and less susceptible to damaging UV radiation (Hartmann, 1996). While more research is needed to establish a definitive link between metabolite levels and function *in planta*, our findings reinforce trends seen in other beverage plants and illustrate the importance of considering common ecological theories when trying to predict patterns of metabolite abundance.

While the effect of plant genotype was not the focus of this study, genotype effects were detected, both in the quantified and untargeted data. Notably, leaves collected from the Arkansas genotype contained significantly more theacrine than the other two genotypes when averaged across leaf stages and treatments (Tukey post-hoc adjusted p = 0.01), suggesting there may be natural variation in methylxanthine metabolism within the species that is genetic in nature. Such variants may be useful to future researchers interested in mapping genetic variation in the production of key metabolites.

The roasting process highly elevated a small number of chemicals, which we hypothesize are the products of both enzymatic and non-enzymatic browning processes. Many such chemicals are derived from the heat-catalyzed reaction of precursor molecules like sugars and chlorogenic acids and are responsible for the distinctive sensory features of coffee and roasted teas. Among the compounds found to be highly elevated in roasted yaupon was larixinic acid (maltol), a Maillard reaction product with a sweet, caramel-like taste/odor (Kato, 2003). Diacetyl trimer, a ketal produced during the coffee roasting process associated with buttery sensory notes, was also highly elevated (Blank et al., 1991; Hotchko, 2014). These two compounds are valued not only as aroma and flavor components but as key markers of roast quality and aroma profile in coffee (Angeloni et al., 2021; Parenti et al., 2014; Yang et al., 2016). Future work investigating the effects of variable roasting conditions may benefit from monitoring these compounds when benchmarking roasting techniques. Several chlorogenic acids also went up significantly with roasting, including multiple predicted isomers of caffeoylshikimic and feruloylquinic acid. Caffeoylshikimic acids have been documented in roasted mate and are proposed to be derived from the dehydration of chlorogenic acids (Jaiswal et al., 2010). Feruloylquinic acid is a chlorogenic acid found in coffee, tea, and yerba maté that has antioxidant properties (Bastos et al., 2007; Boulebd et al., 2023; Clifford & Ramirez-Martinez, 1991). Chlorogenic acids in general are known to have complex effects on sensory attributes, including heat-catalyzed degradation into volatile compounds and solubilization of other flavor compounds (King & Solms, 1981; Moon et al., 2009).

Overall, many similarities were found between the compounds produced during the yaupon roasting process and those found in roasted coffee and tea. Given these parallels, future research on yaupon processing may be able to draw from those bodies of literature to make better predictions on what roasting conditions may be optimal for a range of desired product types. As research in yaupon continues to grow, avenues of study will include quantifying these compounds’ absolute concentrations in yaupon tea, associating them with specific sensory profiles, and measuring how their levels change under differing roasting conditions.

Hundreds of compounds beyond the methylxanthines and chlorogenic acid quantified were found to be strongly associated with leaf stage. These age-associated compounds almost certainly belie nutraceutical and sensory shifts in the final product. Therefore, future researchers interested in when during leaf development chemical changes occur may benefit from collecting tissue at more densely spaced timepoints. As more knowledge is gained about these chemicals, their bioactivity, and their effects on flavor, the yaupon tea industry will be able to tailor its harvesting and processing methods to maximize the palatability and nutritional value of its product.

## 5. Conclusions

This study investigated how two variables important in commercial tea production, leaf developmental stage and roasting treatment, affected the metabolic profile of yaupon leaves across three genotypes of distinct geographical origin. Targeted UHPLC-MS analysis showed that methylxanthine (i.e., caffeine, theobromine and theacrine) and chlorogenic acid (3-CQA) content was highest in the youngest leaves and lowest in mature leaves, but that the roasting conditions tested did not significantly affect levels of either 3-CQA or the three methylxanthines. Roasting did, however, significantly elevate 100 compounds tentatively identified through untargeted UHPLC-MS profiling. Many of these have been documented as sensory chemicals in other roasted teas/coffees and are likely the products of Maillard reactions and other browning processes. Principal component analysis further revealed distinct metabolic patterns based on leaf stage at harvest, suggesting its importance as a variable in commercial harvest. Put together, these findings provide valuable information for the yaupon growing and breeding industry and act as a foundation for more detailed studies to explore the nutraceutical benefits of yaupon holly. Coffee, *Camellia* tea, and yerba mate have all benefitted from combining metabolomics with grower/processor knowledge and intuition. As the budding yaupon tea industry finds its footing, the use of tools like UHPLC-MS promise an exciting future for this long-overlooked American tea.

## CRediT Author Statement

**Ben Long:** Conceptualization, Methodology, Investigation, Formal Analysis, Resources, Writing - Original Draft, Visualization **Scott Harding:** Conceptualization, Methodology, Resources, Writing – Review and Editing **Khadijeh Mozaffari:** Methodology, Software, Formal Analysis, Writing – Review and Editing **Jeffrey L. Bennetzen:** Supervision, Project administration, Funding acquisition, Writing – Review and Editing.

---

**Figure S1.**
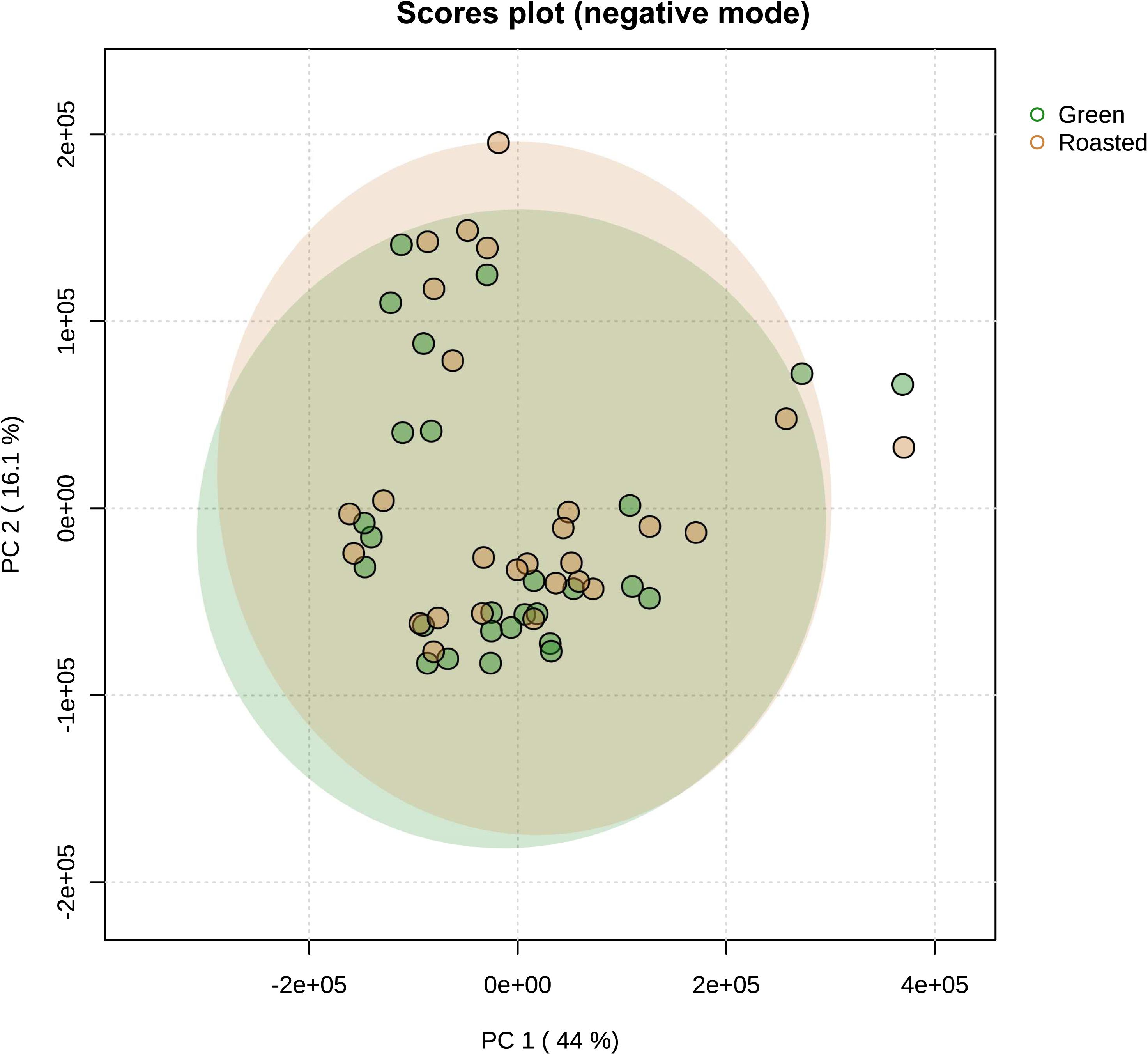
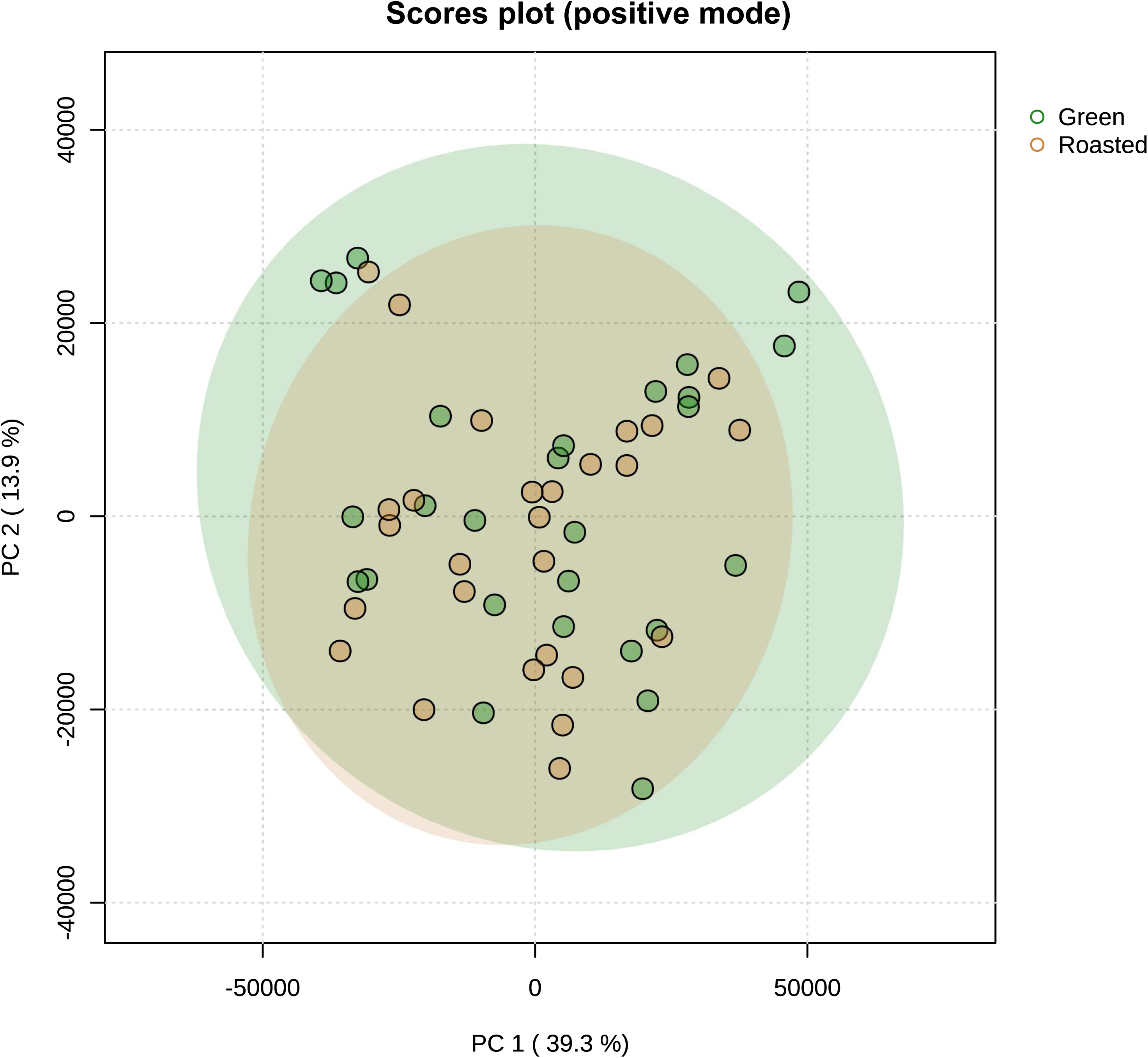
PCA scores plot of samples grouped by processing method. (A) Negative mode results, (B) Positive mode results.

**Figure S2.**
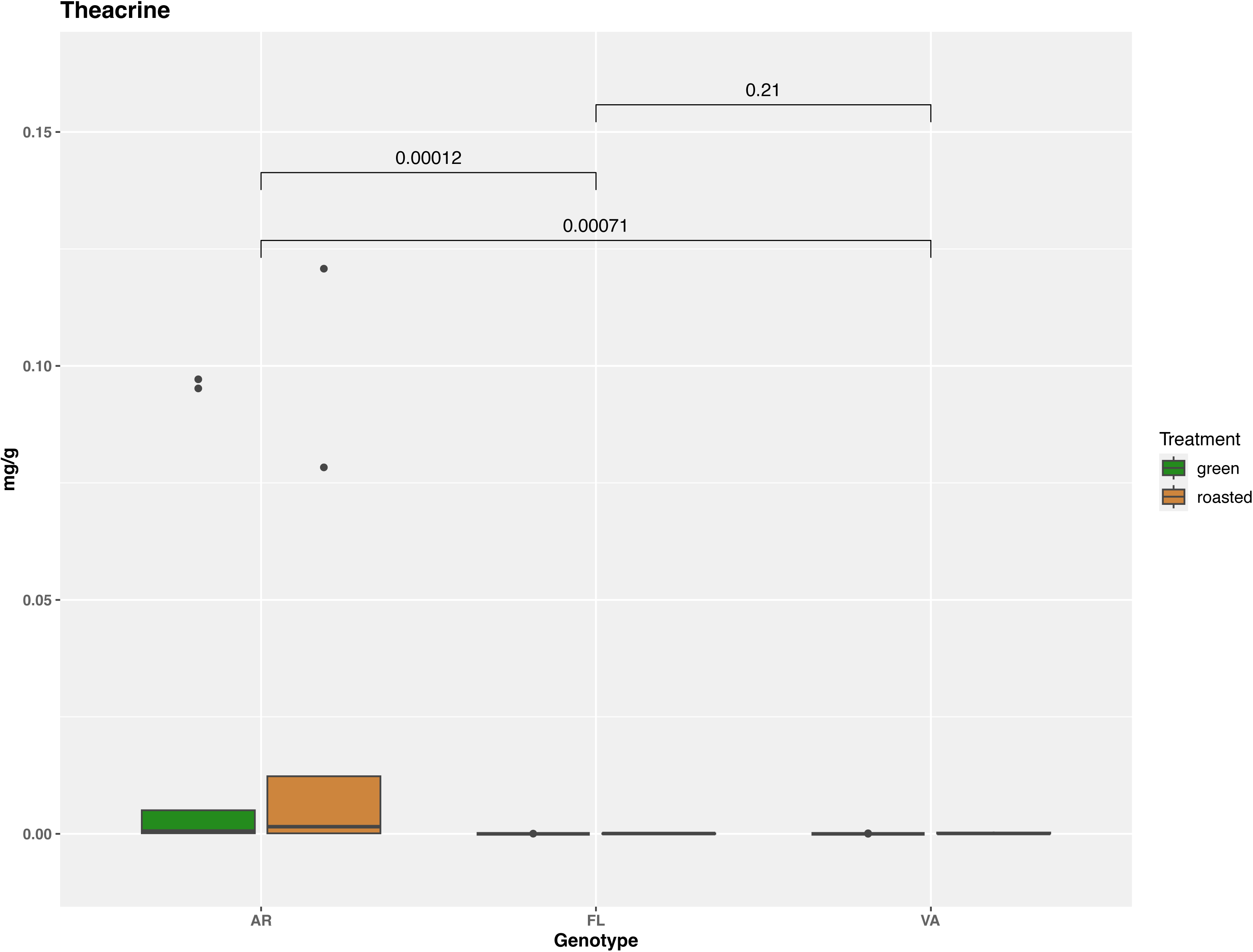
Theacrine content of yaupon holly leaves across all leaf stages and treatments, grouped by genotype.

**Table S1. List of identified chemicals detected in negative ion mode significantly differentiated by leaf stage at harvest.**

**Table S2. List of identified chemicals detected in positive ion mode significantly differentiated by leaf stage at harvest.**

**Table S3. List of identified chemicals detected in negative ion mode significantly differentiated by processing method.**

**Table S4. List of identified chemicals detected in positive ion mode significantly differentiated by processing method.**

